# Hand Drawing Image based Causal Representation Learning for Robust Parkinson’s Disease Feature Extraction and Detection

**DOI:** 10.1101/2025.06.01.657220

**Authors:** Kim Dae young

## Abstract

Being an irreversible disorder regarding the human motor-system, Parkinson’s Disease(PD) has been a threat to many neurological patients, especially due to its severity in pain and muscle control restriction. As PD has no significant cure or treatments to this day, early diagnosis, or detections of PD within potential patients is a crucial task to maximize the effect of mediations which are implemented to achieve temporal prohibition of motor failure progression. In recent research, alongside conventional diagnosis methods based on neurological examinations or MRI based brain imaging, use of deep learning based artificial intelligence models, such as ResNet, are repeatedly reported to have significant progress in detecting PD in early stages with high performance. Based on current success, this research attempted to further enhance AI-driven PD diagnosis by developing a deep learning based causal representation learning framework that extracts only highly robust PD features from simple hand drawings. Specifically, convolutional VAE based reconstruction and information theory based weakly supervised learning were linked with causal representation learning methods to distinguish significant PD features from geometrical features within hand drawing images. Not only aiding conventional tests for PD diagnosis, but also giving reliable representations of PD features such as tremor and rigidity, developed framework was found to achieve high performance in both retrieving latent factors of PD in images and predicting PD diagnosis results.

## I. Introduction

To this day, Parkinson’s Disease(PD) is considered as one of the most crucial neurological damage disorders along with dementia and Alzheimer’s Disease. Though most medical researches focus on finding effective treatments or pathologies to cure a specific disease, irreversible progressiveness in the deterioration of human-motor systems, which can also lead to severe mental breakdowns, and the absence of effective treatments have made PD related researches to focus more on probing for enhanced measures that can allow earlier detections of PD developments with validity [1–3]. Conventional Methods to detect PD highly relied on human-based interpretations. Constructing neurological examinations through assignments, such as Functional Gait Assessment(FGA) tests or geometrical hand drawings, or obtaining brain images through MRI scans, interpretation of data based on domain knowledge was mainly driven by human experts to diagnose PD existence.

However, due to the lack of PD specialists in low and middle income countries (LMIC) and the continuous prevalence of PD in global aspects, there still exist limitations in fully achieving proactive diagnosis and mediations in PD [1,2]. Deep learning can alleviate this burden by assisting experts with valid detection or extractions of PD features. For example, applications of Convolutional Neural Networks (CNN) or Machine Learning algorithms such as RandomForest or Logistic Regression classifiers in detecting PD from voice recording or speech-based spectrogram images demonstrated highly significant classification performance in various research [4–7]. Furthermore, in hand drawing evaluation tasks, shallow CNNs or advanced models such as LeNet led to successful detections of PD symptoms based on images and sensor data from geometrical drawings, such as pen pressure, tilt status, or acceleration [8–10]. Based on current success, and to further enhance earlier PD diagnosis under non-ideal hand drawing environments, such as absence of sensor data or timestamps, this research attempted to develop a hand drawing based causal PD detection algorithm which only requires partial image of geometrical lines drawn by participants. Unlike conventional hand drawing examinations, which required timestamp data, drawing speed data, baseline diagram data, pen pressure and human expert-based evaluation for PD diagnosis, the proposed framework solely implements completed drawings from individuals. Under weakly supervised causal representation learning and VAE based inpainting procedures, extremely robust causal factors that correspond with valid PD diagnosis were extracted and applied to downstream tasks to enhance performance of machine-based PD detections. Results show that the proposed framework can find robust, task-invariant PD features with significance, which can lead to high versatility in diverse drawing evaluation tasks.

## II. Preliminaries

### A. Current Parkinson’s Disease Diagnosis Mechanism

Though no specific cure is found, diagnosis of Parkinson’s Disease can be conducted under clearly defined features, which are mainly related to damaged motor-systems. Among motor-related features, tremor, bradykinesia, and rigidity are considered as primary symptoms of PD [1,2,3,12,13]. Tremor due to PD, also known as shaking, is represented by rhythmic motions, which mostly begin in hands of patients. Proportional to the intensity of stress or intention of movements, tremor can be often characterized by rubbing fingers and tends to be alleviated during sleep. Meanwhile, Bradykinesia, also known as slowness in motions, is another major feature of PD which is characterized by significant decrease in the velocity of voluntary movements [11]. In previous research, bradykinesia is known to be affected by both rest and action tremors in PD patients [12]. Lastly, rigidity, also known as stiffness in motor-systems, is also an important symptom of PD. Being prevalent in most PD patients and known to affect the intensity of tremor or even potentially bradykinesia [12,13], rigidity restricts motions and relaxation of muscles, leading to short or abrupt (non-smooth) movements.

Among PD detection examinations, motor-based features of PD can be effectively detected under hand drawing tasks [14]. As trace of pens, deviations from baseline routes, and acceleration or deceleration of motions can be visualized in hand drawings, obtained images of geometrical drawings can efficiently reflect priorly discussed motor-damage related features, representing existence of PD among participants. Hence, causal links in PD diagnosis can be simplified to a causal DAG as in Fig 1. Based on the fact that high level features can be extracted from low level data using causal representation learning, this study attempted to find key motor-related PD features for robust detection of patients under Fig 1.

**Fig 1.**
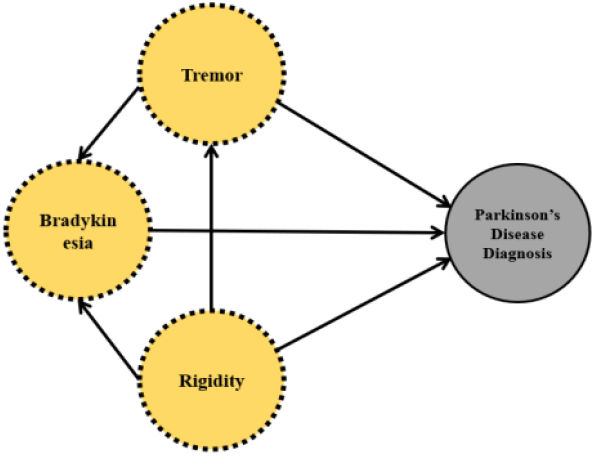
Simplified Causal PD Diagnosis Mechanism.

### B. Related Works

Regarding ML/DL applications in Parkinson’s Disease analysis, basic CNN models or advanced convolutional architectures such as VGG or ResNet were broadly used in most research. In researches which attempted to detect PD in voice data from patients, CNN showed significantly better classification accuracies when compared to conventional ML methods. For example, in [4], pretrained InceptionV3 CNN was implemented to process spectrogram images from telephonic voice datasets: UAMS and mPower to diagnose PD, leading to 97%, 94% accuracies in UAMS/mPower mel-scale spectrogram image classification tasks. ResNet50 and Xception architecture based multiple-finetuned CNNs were also implemented to detect PD from voice recordings [5]. Using vowel based short-duration voice data such as of PC-GITA datasets [7], ensembles from two models each succeeded in achieving 91.33% accuracy and 91% accuracy regarding PCGITA/u/ segment data after enhancing spectrogram images using gaussian blurring. In [6], obtaining denoised spectrogram images under Variational Mode Decomposition(VMD) and connecting ResNet-18, ResNet-50, ResNet-101 based feature extraction with bi-LSTM models for PD classification based on PC-GITA data, use of CNN was again found effective under 95.34%~98.61% PD classification accuracy. Apart from voice recording based PD detection works, PD diagnosis based on the application of CNNs in hand drawings of geometrical lines was also vastly implemented in various research. For example, electronic pen time series data-based circle/spiral/meander drawings were implemented in [8] to train separate CNNs, which were later combined for harmonic weighted voting-based PD classification tasks. This approach led to an average overall accuracy 78.26% in classifying spiral drawings based on non-ensemble CNN fittings, while an overall accuracy of 93.42%~95.74% was achieved based on voting, which regards all types of drawings. In [9], time series data regarding X,Y coordinates, angles, pressure values to the tablet for each timestamps were acquired from drawing Archimedean spiral pentagons or wire cubes on Wacom tablets, and were classified via a basic CNN with 6 convolutional layers and 3 max-pooling layers. Results show that under data augmentation for balancing labels, the best CNN model achieved 0.9 Kappa, 95.02% accuracy, and 98% precision in detecting PD patients. Meanwhile, comparing performance between different CNN architectures in PD drawing classification was also computed in [10]. Among ImageNet, CIFAR-10, LeNet architectures, ImageNet structure-based CNN achieved 87.14% and 77.53% PD classification accuracy in processing meander and spiral drawings. Furthermore, PD detection utilizing only a small set of wave and spiral geometric line drawings from [14] were also widely attempted in the field of CNN based PD classification research. For example, using fine-tuning in VGG-19 models for PD drawing classification enhancement was implemented in [15], using data from [14]. Results show that using VGG-19 models under 10-fold cross validation, 88.5%(spiral) and 88%(wave) classification accuracy and 92.2%(spiral) and 87.9%(wave) specificity can be achieved with significance. In [16], VGG, ResNet, and Vision Transformer (ViT) with patch size 16 × 16 were applied to the identical data of [14] to check PD detection performance. Results show that, with no augmentation methods such as AugMix or PixMix, ResNet101’s model performance was highest in spiral drawing classification tasks with accuracy of 79.33%, whereas VGG19’s performance was highest in wave drawing classification tasks with accuracy of 93.33%.

Based on related works, this research focused on the fact that causal approaches have not been actively implemented for PD diagnosis from hand drawings. For intensive comparisons with previous CNN based works, this research implemented the spiral/wave dataset from [14] within analysis. Checking performance in downstream tasks by implementing latent causal representations extracted from analysis, reliable, yet effective deep learning frameworks for robust PD diagnosis were pursued throughout the overall process.

### C. Causal Representation Learning

Based on the assumption in II.A., to extract robust PD features from hand drawings, causal representation learning was implemented within the proposed framework. Causal representation learning(CRL) is a technique to probe for representational spaces where individual factors are linked under causal relationships. Specifically, deep learning based CRL approach incorporates Neural Network(NN) frameworks for causal factor extractions [17]. Under the assumption of Pearl, J’s structural causal model(SCM) [18,19] (1), DL-based CRL extracts causal information such as direct acyclic causal graphs (causal DAG) using NN approximation, assuming an expanded causal structure from (1) to (2). Unlike conventional ML/DL approaches, which can only incorporate correlation structures among variables, causal representation learning succeeds in extracting complex causal links under SCM assumptions with only gradient descent algorithmic approaches.

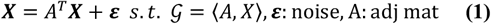

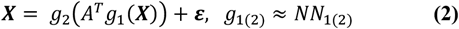

In recent works, probing for causal mechanisms in image data is becoming prevalent to achieve higher robustness and reliability in machine-based decisions [20,21]. To extract causal structures from image data, Latent Causal Representation Learning, which is defined as extracting high level features from low level features, such as pixels, becomes a key process. Based on a generative process between observable variables *X* and latent causal factors *Z* in (3) to (5), both generation function *g* and causal adjacency matrix *A* are learned to find latent causal relationships using generative deep learning frameworks, such as diffusion models, GAN, or VAE [20–23]. In current research fields, methodologies of latent causal representation learning vary by the usage of pairwise inputs, or the usage of error terms in latent space for implicit learning [21,24,25]. For example, [21] implemented both a pairwise input approach and an implicit learning approach under weak supervision to extract latent causal factors such as color of lights or location of objects in general images. Meanwhile, in [25], a non-pairwise approach with the usage of error terms was implemented for causal factor extractions such as light position and shadow lengths in pendulum datasets. Among various methodologies, this research implemented a non-pairwise approach for realistic approaches, where factor-wise pairs cannot be acquired, and an explicit learning approach with Graph Autoencoders which directly encodes latent causal factors instead of error terms to make the overall process as simple as possible.

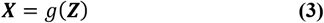

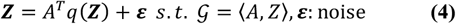

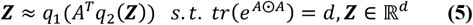

## III. Methodology

### A. Data

Hand drawing image dataset from Parkinson’s disease patients and healthy control group participants were acquired from [14], which is accessible through Kaggle. Based on UPDRS scores, 55 right hand dominant volunteers were divided into two groups: PD patient group with 27 participants, and an UPDRS = 0 control group with 28 participants. Among 27 PD patients, 12 were classified as severity 1 group with modified H&Y of 1 and 1.5, 8 were classified as severity 2 group with modified H&Y of 2 and 2.5, and 7 were classified as severity 3 group with 3 or more modified H&Y scales.

Acquired hand drawing dataset, which was publicly published in Kaggle Datasets, contains 102 spiral hand drawings and 102 wave hand drawings from participants. Each sub-dataset was pre-divided into training sets and test sets, each containing (36, 36) cases and (15, 15) cases of (PD, non-PD) hand drawings. This study implemented both spiral and wave training images(size: 224 × 224, 144 samples) for causal representation learning, and implemented all testing images(60 images) to check downstream task performance. Unlike other works which compute data augmentation to both increase dataset size and robustness of fitted models, this research did not artificially increased size of data to consider small sample based representation learning. Instead, to increase robustness of fitting, image inpainting frameworks were applied for each image of drawings. That is, each sample’s center region was masked with 0 based square patches(size: 100 × 100) as in Fig 2., and was in-painted in the model fitting process. Using inpainting when retrieving latent variables, extremely robust causal representations of factors dealt in Fig 1. were probed for PD detection.

**Fig 2.**
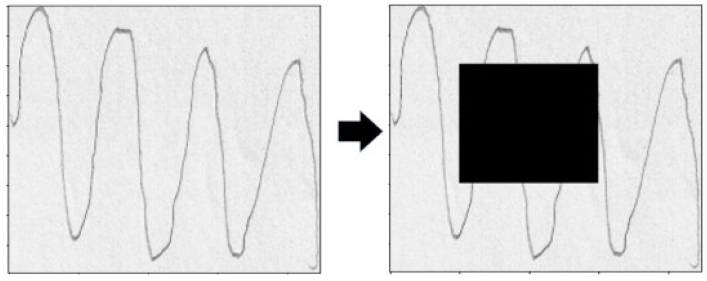
Masking Procedure Example. Left: original wave image. Right: masked image with 100 × 100 size patch.

### B. Latent causal representation learning from drawings

Using 144 images of spiral/wave hand drawings as input, latent causal representation learning was computed with a novel framework depicted in Fig 3. For latent space extraction, Variational Auto-Encoder(VAE) [26] was implemented with *L* set as 1(sampling once) and a gaussian normal distribution set for random noise within reparameterization. For the encoder structure of VAE, three convolutional layers, which were set as having 64 4×4 filters with stride 2, 32 3×3 filters with stride 3, and 16 3×3 filters with stride 2, sequentially, and two fully connected(FC) layers with 1024 and 128 hidden nodes after flattening were implemented. Meanwhile, regarding latent space dimensions, 4 latent variables were assumed to extract three causal factors that correspond with features in Fig 1., and one geometrical factor that represents geometrical features from spiral or wave itself. For decoder in VAE, encoder-symmetric structures with two FC layers and transposed convolutional layers were implemented for image reconstruction.

**Fig 3.**
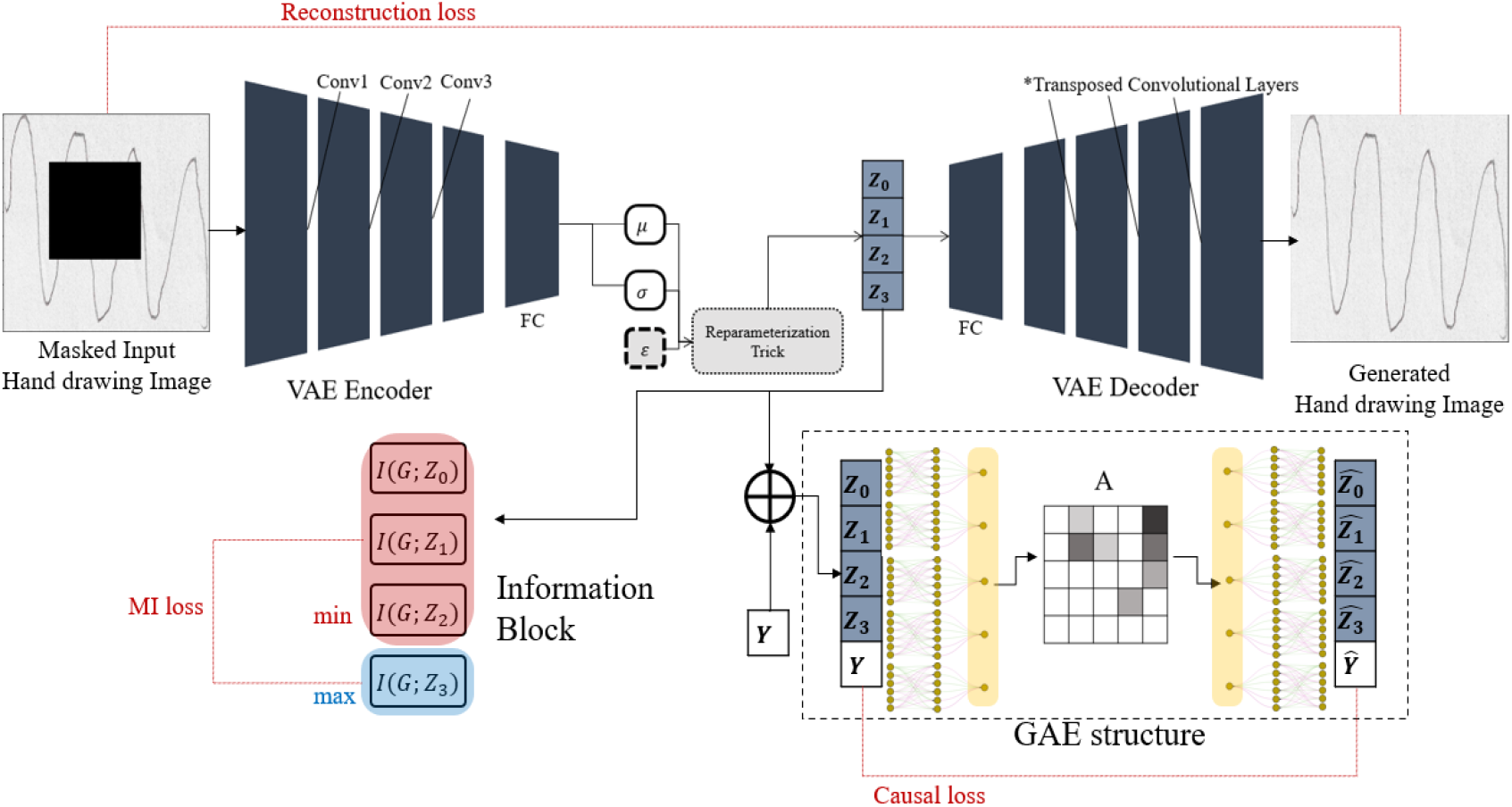
Visualization of Latent Causal Representation Learning from PD Hand Drawings (VAE+GAE+MI approach).

**Fig 3.**
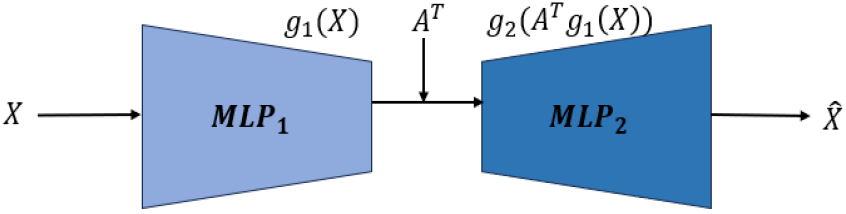
Basic GAE framework. *A*: adjacency matrix

Meanwhile, causal representation learning in latent space was computed using the Graph Autoencoder(GAE) approach explained in III.C. Here, to apply weakly supervised learning, concatenating target variable (Parkinson Disease diagnosis: 0(healthy), 1(PD)) to the extracted latent vector *Z* was computed for GAE inputs. In addition, for variables: *Z*_0_, *Z*_1_, *Z*_2_ to focus only on PD related factors, geometrical-task related features were reinforced to be encoded to latent variable *Z*_3_ by incorporating a Mutual Information(MI) approach. Adding an Information Block which calculates the MI between *Z*_*i*_ and auxiliary binary variables *G* (spiral: 0, wave: 1), weights in VAE were updated to maximize *I*(*Z*_3_;*G*) and minimize *I*(*Z*_*i*_;*G*) (i=0,1,2) under loss function ℒ_*IB*_. Loss function for training weights was computed as (6). Total Loss was computed based on the sum of individual Loss functions of VAE, GAE, and the Information Block. For VAE loss, Evidence Lower Bound (ELBO) of (7) was implemented to approximate MLE approach. Assuming identity covariance matrix, ELBO was further reduced into the Mean Squared Error form in (8). Though different weights for the in-painted area and non-masked area can be assumed for MSE, as this research focuses on extracting latent causal features, region-based weighting was not applied when computing MSE for reconstructed hand drawing images.

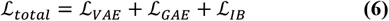

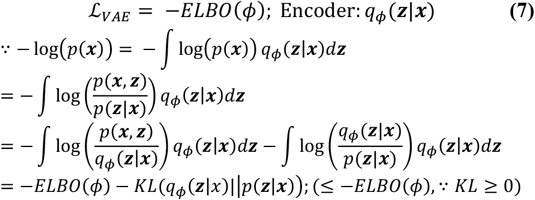

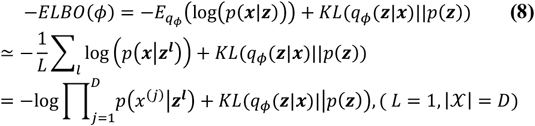

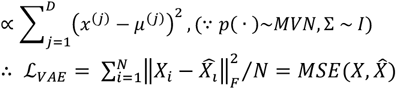

For the Information Block loss, negative value of MI: *I*(*Z*_3_;*G*), mean value of *I*(*Z*_*i*_;*G*) (i=0,1,2), and weight parameter *v* were implemented as (9), considering the overall minimization objective of optimization. Thus, ℒ_*total*_ was comprised of two MSE results from VAE and GAE, and a weighted Mutual Information statistic between latent variables and auxiliary variables *G*. The overall optimization process for PD hand drawing based latent CRL was summarized as **Algorithm 1**.

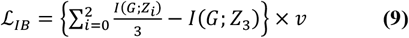

Mutual Information *I*(*G*; *Z*_3_) = *H*(*G*) + *H*(*Z*_3_) − *H*(*G, Z*_3_)

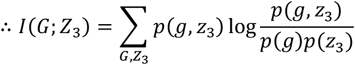

#### Algorithm 1

Optimization Process for Latent CRL

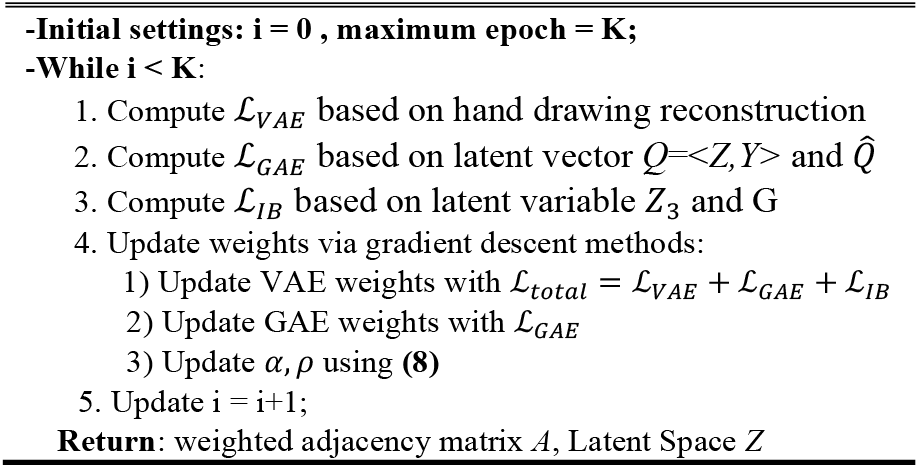

### C. Graph Autoencoder Structures

Within CRL approaches, valid settings regarding DAG extraction methods become crucial. As it was proven by Xia, K et al.(2022) that DNNs with causal graph 𝒢 as a constraint can achieve full expressivity and canonical causal inference under the theorem of 𝒢-consistency and *L*_2_ - 𝒢 representation [27], valid extraction methods that can return causal graph 𝒢 become both a prerequisite and a sufficient condition for neural causal representation learning. Regarding recent causal graph extraction methods, especially DAGs, three algorithms: NOTEARs, Graph Autoencoder, and DAG-GNN [28–30] can be considered as representative works that demonstrate high validity and applicability. This research implemented the Graph Autoencoder(GAE) approach for analysis [29], assuming PD features as causal factors. Under SCM assumptions as in (2) and (5), and incorporating the idea of message passing in Graph Convolutional layers, GAE extracts DAG from tabular data based on the framework of Fig 3. Implementing two MLPs for the encoder-decoder structure, and setting a weighted adjacency matrix layer between MLPs, causal relationships between variable set *X* are extracted from training GAE. Hence, working as a high-dimensional function which can incorporate both linearity and non-linearity, MLPs *g*_1_ and *g*_2_ approximate functions in (2), which is a generalized version of SCM. Here, under universal approximation theorem of DNNs, trained GAE’s weighted adjacency *A* was proven to capture underlying causal DAGs with validity [29] under the objective of (10). In (10), *θ*_1(2)_ corresponds to weights in MLPs, *λ* term corresponds to L1 regularization, and *tr*(*e*^*A*⨀*A*^) = *d* denotes the equality constraint for DAG assumptions in *A* [29]. Using Lagrange multiplier of *α*, (10) can be converted into a relaxed form as (11) and (12). Using this approach, weighted adjacency matrix can be easily retrieved with general NN gradient descent algorithms.

In this research, for analytic flexibility, individual MLPs(two hidden layers, 6 nodes for each hidden layer) were assumed for each variable, leading to a general additive approach for latent variables in set *Q* (Fig 4.). Here, weight masking techniques were implemented with TensorFlow’s tf.keras.constraints. Constraint() to embody both the additive structure and the weighted adjacency matrix within GAE while enhancing computational efficiency. In iterative training, specific hyper parameter settings were set as in *D. Experimental Design*.

**Fig 4.**
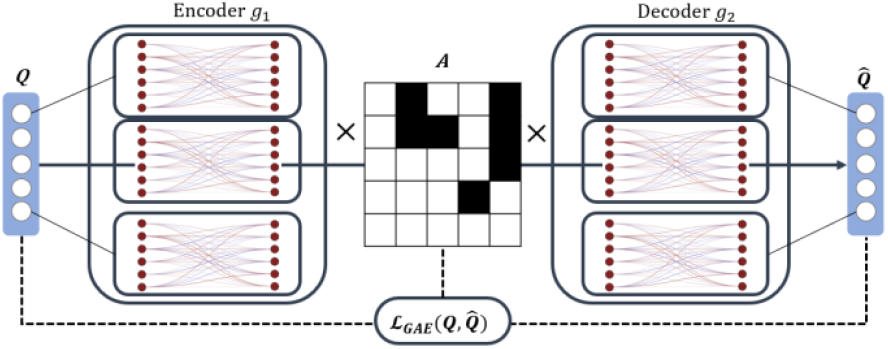
Additive GAE framework visualization. Q denotes concatenated version of latent variable *Z* and target variable *Y*

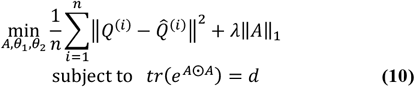

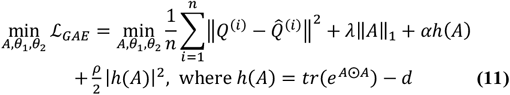

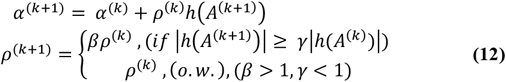

### D. Experimental Design

Latent causal representation learning from hand drawing images and checking causal factor retrievals were processed through training. All model constructions and training procedures were based on Python Tensorflow version 2.18.0. In training the GAE+IT block-based VAE model, different loss functions and learning rates were applied for VAE and GAE weights. For VAE, total loss of ℒ_*Total*_ and an Adam optimizer with *η* = 0.0015 were implemented to incorporate both the image generation objective and the causal representation extraction objective. On the other hand, loss function of ℒ_*GAE*_ and an Adam optimizer with *η* =0.005 were implemented for weights in GAE to only focus on extracting valid adjacency matrix *A*. Furthermore, to only focus on causal links, reconstruction error of *Z*_3_ was not considered when computing MSE in ℒ_*GAE*_. For hyper parameter settings in GAE and the Information Block, *γ, β, λ, v* were set as 0.9, 1.01, 2.0, and 1.0. Initial values of parameters *α* and *ρ* were set as *α*=0.6, *ρ*=0.1. For activation functions, ReLU and ELU were implemented in VAE structures, whereas linear activation functions were implemented in GAE structures. Apart from basic settings, for fast convergence in training weighted adjacency matrix *A*, diagonal elements, outgoing edges from *Y*, and outgoing edges from *Z*_3_ to (*Z*_0_, *Z*_1_, *Z*_2_,*Y*) were blacklisted before fitting. Using **Algorithm 1** based weight updating, 400 epochs were computed under a full-batch gradient descent approach.

When computing Mutual Information values: *I*(*Z*_*i*_; *G*), mutual_info_classif() in python’s sklearn.feature_selection package was implemented with the number of neighbors set as 5. After training and convergence check being completed, fitted weighted adjacency matrix *A* was binarized by setting the overall mean of absolute weights in matrix *A* as the threshold value. Values over the threshold were converted to 1, whereas values below the threshold were converted to 0. Using the binarized adjacency matrix of *A*, latent causal graph was extracted and was compared with the domain causal DAG proposed in Fig 1. Using best match within isomorphic graphs, causal retrieval was evaluated by calculating the Structural Hamming Distance (SHD) between two graphs. Specifically, as this research focuses on causal links related to PD, the 2-hop sub-graph from target variable *Y* (PD diagnosis) was mainly compared with the domain causal DAG. Lastly, for intense validation of latent space from GAE+IT block-based VAE, disentanglement was checked using both quantitative, qualitative measures. First, for geometrical representation *Z*_3_, quantitative evaluations were mainly computed. By setting latent variables in *Z* as code dimensions and setting geometric labels *G* and PD labels *Y* as factors, modularity score (disentanglement score) in [32] was computed based on LASSO regressors as introduced in [31]. Specific formula of modularity follows definitions in [31,32]: (13)-(14).

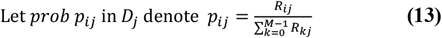

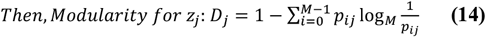

Here, the term’ *R*_*ij*_ ‘denotes the magnitude of the weight (coefficient) which corresponds to code dimension *z*_*j*_ when target variable(factor) is set as *v*_*i*_ in regression tasks. In this work, number of factors was set as 2(*Y, G*) and the number of code dimensions was set as 4(dimension of latent space), when computing modularity. For LASSO regression implementations, Python’s sklearn.linear_model package’s Lasso() function with

*α* set as 0.1 was used for disentanglement score computations. Second, for causal features regarding PD: *Z*_0_ to *Z*_2_, as no specific multi-label data was given for tremor, bradykinesia, or rigidity, implementing quantitative measures under predictors had a high possibility of underestimating success in practical disentanglement achievements. Thus, for *Z*_0_ to *Z*_2_, visual comparisons based on varying individual latent variable values were mainly driven for disentanglement checks. By varying each variable with other latent space being fixed, changes in the generated image were qualitatively evaluated to check clear existence of visual characteristics which correspond with representative features of tremor, rigidity, or bradykinesia. (*All experiments and model constructions were processed using Google Colab’s basic T4 NVIDIA GPU environments).

### E. Parkinson’s Disease Prediction using hand drawings

Based on success in latent causal representation learning from hand drawing images for PD detection, the feasibility of using extracted robust latent spaces in downstream tasks was evaluated under Machine Learning based predictions. Using latent representations retrieved from the GAE+IT block-based VAE model, train data images were first converted to 4dimensional vectors *Z* to construct new training data. Among latent variables in *Z*, only causal factors: *Z*_0_, *Z*_1_, *Z*_2_ were used to check the effectiveness of implementing causal frameworks in PD detection tasks. Here, to consider intrinsic randomness caused by reparameterization in VAE structures, training data images were converted to 3-dim vectors multiple times with different random states from tf,keras.utils.set_random_seed, leading to an up-sampling effect of hand drawing data in training. Specifically, using 10 different seeds, the initial training dataset of size: (144, 224, 224,3) was converted into a new tabular dataset of size: (1440, 3).

Based on the new train dataset, two shallow DNNs: M1, M2 with 3 hidden layers were fitted for Parkinson’s disease detection using hand drawings from individuals. After training, fitted DNNs were tested under test data hand drawings from [14]. To return stable predictions, when compressing test data images to tabular data with the encoder, the mean value of 10 different latent representations(*10 different random states implemented) from test images were used as input for PD prediction. To check the causal representation + DNN based PD detection performance or prediction performance, classification metrics: accuracy, precision, recall, and f1-scores were implemented. Computing four evaluation metrics for both spiral and wave drawings each, performance of GAE+IT based VAE model usage was compared with existing advanced CNN or Vision Transformer based results, which implemented identical dataset [15,16]. (*In DNN modeling, python tensorflow.keras package was used for skeleton construction. For activation functions, ELU function was set for hidden states, whereas a sigmoid function was implemented for the output layer. For efficiency in training, batch normalization and He-Normal weight initialization was applied to all layers. Further hyper parameter settings were set as in TABLE I.)

**TABLE I.** Hyper parameters settings for DNNs.

| Parameter | Settings |
| --- | --- |
| Hidden states | M1: {16,16,8}, M2: {32,32,16} |
| Optimizer | Adam Optimizer |
| Loss function | Binary Cross Entropy |
| Epochs | 350 |
| Batch size | 200 |
| Learning rate | 5e-3 |

## IV. Experimental results

### A. CVAE and GAE fitting results

Results of fitting GAE+IT block-based VAE were deduced as Fig 5. Within updating neural network weights, convergence in ℒ_*Total*_, ℒ_*VAE*_, ℒ_*GAE*_, ℒ_*IB*_ was checked after 400 epochs of training. For ℒ_*Total*_, stable convergence in values was achieved after 100 epochs in training, resulting in a final loss of 1.6022. For ℒ_*VAE*_, successful convergence to 0 was checked with final MSE being approximately 0.0058. Lastly, for ℒ_*IB*_, convergence was found after 350 epochs, which led to a final value of ℒ_*IB*_= −0.1216. This implies that retrieving valid causal latent space and generating precise hand drawing images while considering inpainting objectives were both successfully achieved under GAE + IT block-based VAE fitting. Specific examples of in-painting based image generations were visualized in Fig 6. Original ground truth images were visualized in the first column, masked original images were visualized in the second column, and reconstructed images were visualized in the third column. Comparing images between ground truth images and generated images, success in fitting encoder-decoder structures in convolutional VAE was also checked in qualitative aspects. In most cases, successful generation of spiral/wave images or valid inpainting was verified in visual aspects. Features such as wavering of lines, darkness due to intensity of pen pressures, and distance between lines were successfully captured and reconstructed from VAE encoder-decoder structures. For example, in the second row in Fig 6., generated image in the third column exhibits highly successful reconstructions of PD related features in the original image, such as collapse of lines within the center region and large deviations from the baseline, leading to a representative example of detecting major motor-related symptoms from nonaugmented spiral images drawn by patients who were diagnosed with PD. (*Comparing spiral and wave hand drawing reconstructions, it was found that generated geometrical lines were more vivid within spiral drawing groups).

**Fig 5.**
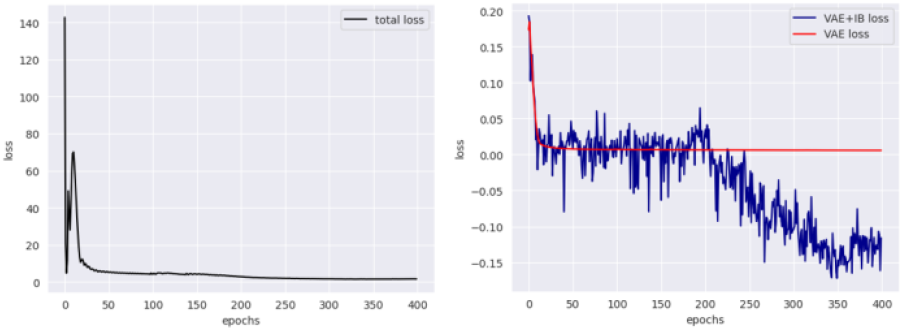
Visualization of loss values from fitting GAE+IT blockbased VAE model. (Left) ℒ_*Total*_ per epoch (Right) ℒ_*VAE*_ +ℒ_*IB*_

**Fig 6.**
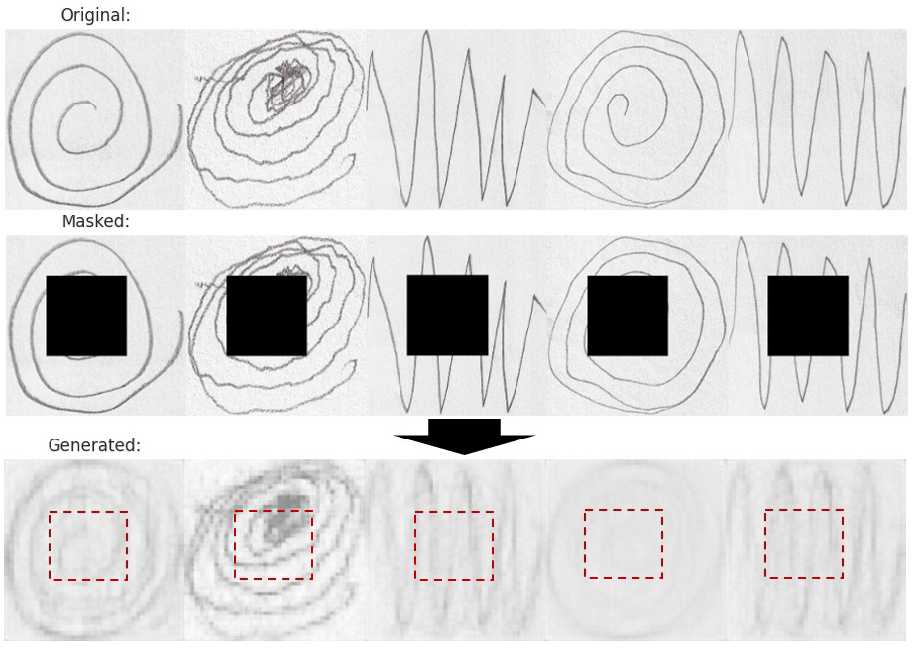
Visualization of generated images under GAE+IT based VAE fitting. First row ~ third row: original hand drawing image, masked corresponding image, and generated image (inpainting completed). Red-dotted box in third row denotes in-painted region.

Under successful fitting and convergence in latent causal space extractions, weighted adjacency matrix between latent variables was returned as Fig 7 (left). Though one edge between *Z*_1_ and *Z*_2_ was found bi-directional, all other causal links extracted in matrix *A* satisfied the acyclicity conditions of DAG. Based on threshold value of 0.0806, weighted adjacency matrix was then binarized as Fig 7 (right). Using binarized adjacency matrix of *A*, 2-hop sub graph from target node was extracted as Fig 8. Based on best match, extracted latent causal sub-graph for PD was compared with the domain Causal DAG from Fig 1. Comparing graphical structures between two graphs, SHD = 2.0 was achieved, which implies that the extracted causal graph and domain causal DAG have significantly high similarity in topological aspects. In addition, as target variable of *Y* was included in both the domain DAG and the extracted graph, it was found reasonable to infer that extracted causal features may have high possibility of representing domain PD factors: tremor, rigidity, and bradykinesia, under one-to-one correspondence.

**Fig 7.**
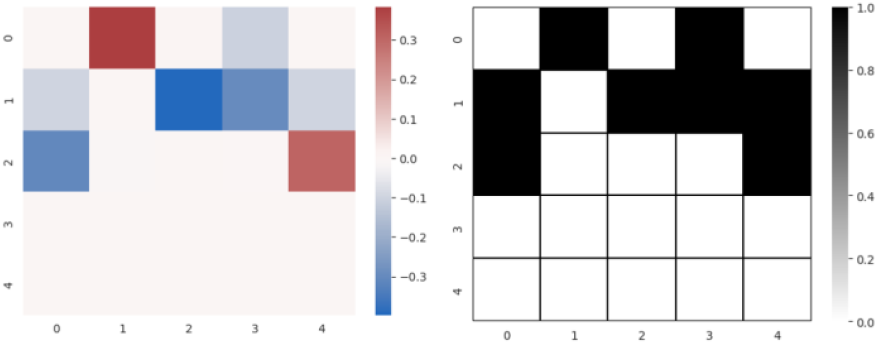
(Left) Latent adjacency matrix (Right) Binarized *A*.

**Fig 8.**
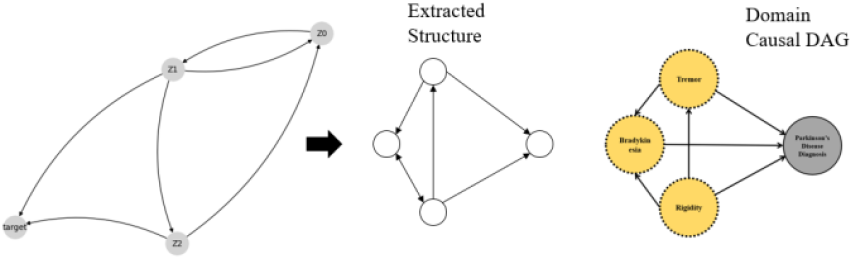
2-hop Subgraph from target node(PD diagnosis).

### B. Causal disentanglement checks

Under successful latent causal graph extraction for Parkinson’s disease, disentanglement regarding variables: *Z*_0_ to *Z*_3_ was checked to validate whether extracted latent variables can be one-to-one matched with domain knowledge-based PD diagnosis factors and geometrical features. First, for *Z*_3_, quantitative evaluation based on the disentanglement score *D*_3_ and qualitative evaluation based on varying only *Z*_3_ values in latent space were both computed for analysis. Regarding *D*_3_ computations, LASSO regression based *p*_*ij*_ computation results were deduced as TABLE II. Based on the definition of (13) and (14), modularity, or disentanglement score for variable *Z*_3_ was computed as *D*_3_ = 0.7094. Having a high disentanglement score close to 1, latent variable *Z*_3_ was found to successfully disentangle geometric features from raw hand drawing images. In other words, this implies that as geometrical baseline features were well separated by latent space of *Z*_3_, remaining latent subspace of variables: *Z*_0_ to *Z*_2_, may have high possibility of containing mostly PD related robust features, regarding the convergence found in ℒ_*Total*_.

**TABLE II.** *p*_*ij*_ MATRIX BASED ON FITTING Lasso regressors under *α* = 0.1.

| $p_{ij}$ values | $v_0(\equiv Y)$ | $v_1(\equiv G)$ |
| --- | --- | --- |
| $Z_0$ | 0.0 | 1.0 |
| $Z_1$ | 0.4306 | 0.5694 |
| $Z_2$ | 0.6226 | 0.3774 |
| $Z_3$ | 0.0510 | 0.9490 |

Second, based on the fact that quantitative measures dealt in *Z*_3_ and TABLE II cannot explicitly represent success in disentanglements for variables *Z*_0_ to *Z*_2_ due to the absence of multi-label datasets which can correspond to major PD symptoms, visual variations were mainly implemented as an alternative to qualitatively assess disentanglement success in latent causal variables. To vary individual latent variables with others fixed, set ={−2,−1.5,−1,−0.5,0,0.5,1,1.5,2}, set= {−3,−2.5,2,−1.5,−1,−0.5,0,0.5,1}, and set={−4.5,−4,−3.5,−3,−2.5,−2,−1.5,−1,0.5} with identical intervals of 0.5 were implemented for possible variations. Changes due to latent variable variations were visualized based on fitted VAEs. Representative disentanglement visualization results from random samples were summarized in Fig 9. Results show that for variables: *Z*_1_, *Z*_2_, and *Z*_3_, successful disentanglement results were found in most visualizations. For example, in varying *Z*_3_, as values increased from −2 to 2, original forms of waves were transformed to spirals with other features being fixed, whereas as values decreased from 2 to −2, original forms of spirals were transformed to waves.

**Fig 9.**
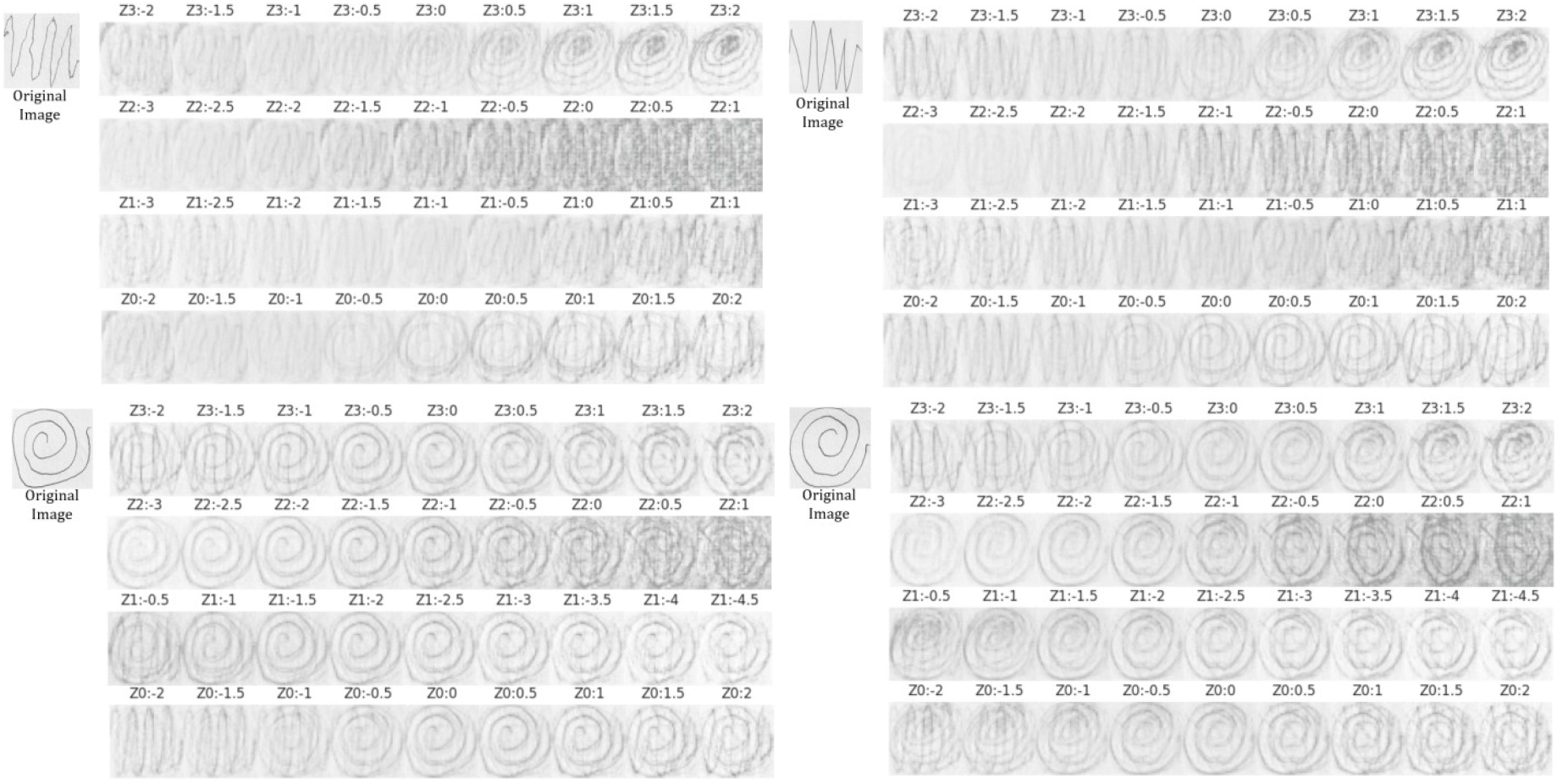
Visualization of Disentanglement checks under individual variations of latent variables: *Z*_3_ to *Z*_0_ (4 random cases: 2 wave images, 2 spiral images). Successful disentanglement results found in *Z*_1_, *Z*_2_, *Z*_3_ variations.

Meanwhile, for causal latent variable *Z*_2_, as values were increased from −3 to 1, only the degree, or intensity of oscillatory errors in geometrical lines were reinforced. Furthermore, in spiral drawing variations, reduction of diameters or increase in compactness with increased intensity of oscillations were visible in visualizations, which is a representative feature of PD related tremor in drawings [33]. Thus, having high commonality in characteristics, latent causal factor *Z*_2_ was not only found to have high possibility of having one-to-one correspondence with tremor in topological aspects, but also found to have one-to-one correspondence with tremors in the aspect of representative features under success in disentanglement.

In *Z*_1_ variation visualization examples, wave hand drawings and spiral hand drawings exhibited different behaviors when *Z*_1_ values were varied with other features set as constant. In disentanglement checks under wave hand drawings, as values of *z*_1_ were increased from −3 to 1, original geometrical lines became cramped, and overall compactness of the wave was increased while other features were fixed. Moreover, reductions in line smoothness were also visible as values of *z*_1_ were increased from −3 to 1, which are all representative features of rigidity in PD [34, 35]. Meanwhile, for disentanglement results under spiral drawings, as values of *z*_1_ increased from −2 to −0.5, original geometrical lines became cramped, and as values *z*_1_ decreased from −2 to −4.5, smoothness of spirals were highly reduced. In other words, deviations from *z*_1_= −2 led to higher prevalence of rigidity-based features with other variables relatively being fixed. Therefore, it was also found valid to say that latent causal factor *Z*_1_ can represent and disentangle PD related symptoms under one-to-one correspondence with rigidity.

On the other hand, unlike latent variables: *Z*_1_ to *Z*_3_, latent causal variable *Z*_0_ exhibited low disentanglement performance in visual aspects, as geometrical baseline features (spiral or wave) itself were found altered or combined in variations. This implies that though the information block succeeded in extracting geometrical baseline features using MI, *Z*_0_ also partially contains information regarding non-PD related features, leading to failure in one-to-one correspondence with the last PD factor, bradykinesia. However, as *Z*_0_ is still a latent causal factor which has causal links with both variable *Z*_1_ and *Z*_2_, it is possible to say that *Z*_0_ might contain other PD related feature information which were not dealt in preliminaries of this research.

Thus, through disentanglement checks, it was found that the proposed GAE+IT based VAE model for PD hand drawing analysis can not only extract significant latent causal spaces but also return precise representations of PD related tremor levels and rigidity levels with high validity.

### C. PD prediction modeling based on latent causal space

Based on success in latent causal DAG retrieval and considering success in disentanglements regarding variables: *Z*_1_ to *Z*_3_, causal variable-based Parkinson’s disease detection modeling was implemented to check performance of extracted latent causal spaces in enhancing success in downstream tasks. Under shallow DNN model structures: M1 and M2, training was computed in Google CoLab environments. After fitting, PD detection performance was checked using test datasets summarized in III.*A*. Among M1 and M2, M1 showed higher performance in PD detections in most metrics. Specific performance metric values for spiral/wave test dataset were summarized in TABLE III (M1). Results show that the proposed latent causal factor-based DNN (M1) can lead to highly significant success in detecting PD from spiral hand drawings. Specifically, in spiral hand drawings, test accuracy, precision, recall, and F1-scores were computed as 90.00%, 92.86%, 86.67%, and 89.66% (Confusion Matrix: Fig 10.). That is, unlike detection performance found in test wave images, which exhibits severe overfitting issues, a significantly light, yet effective PD detection model was achieved in (test databased) spiral image classification tasks with only three latent representations, *Z*_0_ to *Z*_2_. Furthermore, in spiral drawing classifications, 86.67% sensitivity, 93.33% specificity, and 94.67% ROC-AUC score were achieved, which imply reliable predictions can be returned using only three latent causal factors from raw hand drawing images. To further assess success of implementing latent causal representation learning for PD predictions, PD detection performance from spiral images in M1 was compared with other recent works [15,16] which used identical data under advanced DL approaches: TABLE IV.

**Fig 10.**
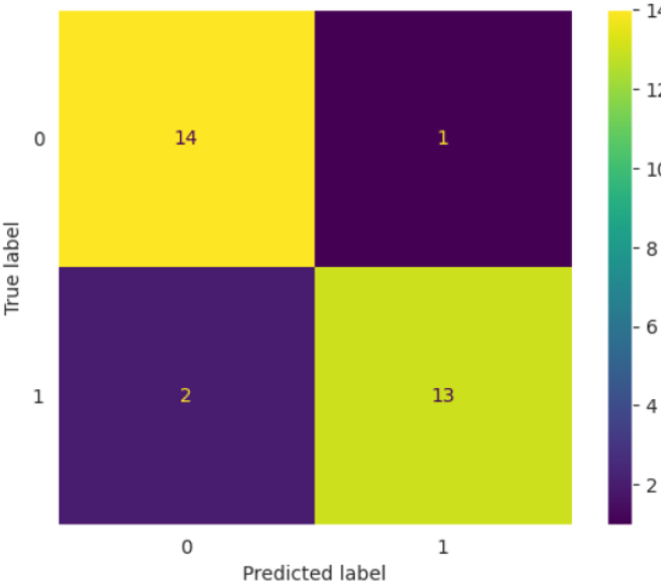
Visualization of Confusion Matrix computed in Spiral hand drawing test dataset classification tasks.

**TABLE III.** Parkinson’s disease detection performance: M1.

| Train/Test | Accuracy | Precision | Recall | F1-score |
| --- | --- | --- | --- | --- |
| Spiral(Tr) | 0.8944 | 0.9152 | 0.8694 | 0.8917 |
| Spiral(Te) | **0.9000** | **0.9286** | **0.8667** | **0.8966** |
| Wave(Tr) | 0.9417 | 0.9141 | 0.9750 | 0.9435 |
| Wave(Te) | 0.5333 | 0.5455 | 0.4000 | 0.4615 |

**TABLE IV.**
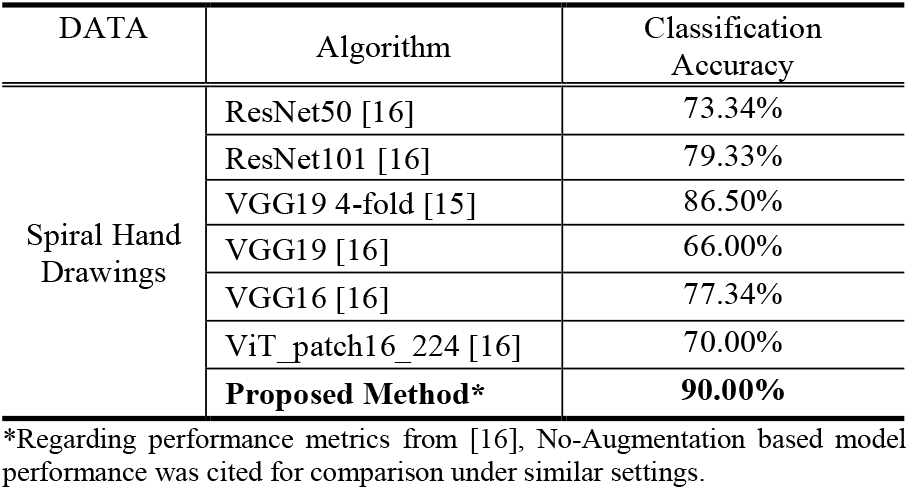
Parkinson’s disease detection performance comparison(spiral)

Under comparison, it was found that the proposed methodology in this research outperforms advanced CNN architecture-based or ViT-based approaches in all cases in the aspect of classification accuracy. For example, the causal approach in this work exhibited higher classification accuracy when compared to (no-augmentation based) Vision Transformers with 16 × 16 sized patches, VGG-19(4-fold), VGG-16, and ResNet101 models [15, 16], by having 20.00%p, 3.50%p, 12.66%p and 10.67%p higher classification accuracy. That is, the proposed framework in this research exhibits high competitiveness and sufficient potential compared to existing methods, though only three causal representation factors from spiral hand drawing images were used as input for simple DNN structures with no specific optimization of hyper parameters. Hence, it was found that latent causal representation learning based approaches and variables computed from this framework can extract robust representation of features directly related to PD diagnosis, while leading to successful performance within downstream tasks: raw spiral hand drawing image-based PD detection.

## V. Conclusion

In this study, latent causal representation learning was implemented to extract robust features of Parkinson’s Disease from hand drawing images. Using generative modelling under VAE structures and implementing Graph Autoencoder-based adjacency matrix learning with weak supervision, a valid causal graph which corresponds with the domain-knowledge based PD diagnosis causal DAG was successfully extracted. Furthermore, under quantitative and qualitative disentanglement checks, it was found that extracted latent causal factors can one-to-one correspond with major PD features, such as tremor and rigidity. Successfully capturing causal mechanisms of PD diagnosis, valid simulations, or successful estimations of treatment effects regarding severity of PD can be computed through the extracted causal mechanism when certain interventions for specific symptoms are assumed. Based on this framework, this research also evaluated the effectiveness and validity of using causal approaches in PD detection tasks with only images of hand drawings. Results show that in spiral image processing tasks, high performance(Accuracy=90.00%) can be achieved by connecting DNN based models with only three causal representations: *Z*_0_ to *Z*_2_. Under competitiveness found in comparisons with advanced CNN architectures such as VGG16, ResNet101 or Vision Transformers, using latent causal space was found to be effective in both extremely decreasing the number of parameters in current image processing models and enhancing robustness in classifications for Parkinson’s disease detections.

Although significant success was achieved in multiple aspects, certain limitations still remain such as eliminating over-fitting issues found in PD detection performance results under wave based drawings environments, or finding exact latent representations that can one-to-one correspond with bradykinesia. For further enhancements, fine-tuning or concatenating non-causal information in downstream task models or implementations of more advanced causal mechanisms to extract a broader range of PD related features should be implemented for better performance.

